# Virucidal Activity of Potassium Hydroxide Modeling on Tospovirus

**DOI:** 10.1101/2022.09.12.507442

**Authors:** Emin Zumrutdal, Muharrem Arap Kamberoglu, Havva Nur Saglam

## Abstract

Viral agents that cause disease in the respiratory system have led to widespread health problems in the world. The continuation of mutations in these viruses and the lack of an effective treatment agent bring possible public health risks. In this study, the virucidal activity of potassium hydroxide (KOH) was evaluated. The intermolecular interactions of KOH and envelope-structured molecules in enveloped viruses and the virucidal activity of these interactions on Tospovirus, which has the ability to infect, were evaluated.

For this study model, the intermolecular interactions of KOH in the lipid bilayer of the virus envelope were evaluated in silico by Doking method. Then, the plant virulence ability of Tospovirus was observed by the direct interaction of KOH with Tospovirus.

It was observed that KOH interacted exergonically with the glycerophosphate structure in the envelope structure. It was determined by clinical and laboratory observations that Tospovirus in plants lost its virucidal activity after interaction with KOH.

In the light of this information, it was thought that KOH had a virucidal effect in enveloped viruses. It is thought that KOH creates this virucidal activity by KOH-glycerophosphate intermolecular interactions and viral envelope lipid layer hydrolysis.

The mucolytic, alkalinizing and possible low-weight immunoglobulin-forming potential of KOH has been demonstrated in previous studies and no pathology was detected in toxicity studies in mice.

In the light of this information, optimized KOH inhalation has a very serious potential as a virucidal agent in diseases caused by enveloped viruses in the respiratory system such as Coronavirus, H. influenza.

## INTRODUCTION

Epidemics and pandemics caused by viral diseases cause serious health problems and deaths in the world, and also negatively affect the sustainability of life(1). Epidemics such as Spanish flu caused by H.influenza, SARS, MERS and Covid-19 caused by Coronavirus are serious examples. These viruses, which cause epidemics and pandemics, usually cause diseases in the respiratory tract. The causative agents are often enveloped viruses(2). Hantavirus pulmonary syndrome (HPS) seen in the USA caused by Hanta virus, Hemorrhagic Fever Renal Syndrome (HFRS) seen in Europe and Asia and diseases caused by other enveloped viruses can be added to these(3).

Viruses generally consist of genetic material containing an RNA or DNA chain, a capsomere, and in some viruses, an envelope that encloses this genetic material. In addition to the very small structure of the virus, the very easy mutation of the virus structure makes the studies for the treatment of viral diseases very difficult(4,5).

Some DNA (such as Herpesviruses, Poxviruses) and RNA viruses (such as Arenaviruses, Coronavirus) are released from cells by budding and wrapping in cell membranes as they mature. Therefore, the structure of the virus envelope is very similar to the molecular structure of cell membranes. Although the bilayer envelope is encoded by the host cells, the peplomers (glycoprotein structure) in the envelope are differentiated by viruses. Glycoproteins, also known as specialized peplomers, on the envelope structure increase their infectivity with their virus-specific special protein (such as spike protein, E protein, HE protein, neuraminidase protein, F-protein) structures(5,6).

In enveloped viruses, the cell envelope lipid layer is structurally very similar to the human cell membrane. Although the molecular content of the cell envelope includes a wide variety of molecular structures, lipid structures are very common (7). In this structure, there are separate hydrophilic and hydrophobic parts. These structures are very important for the protection of the genetic materials that enable the reproduction in the virus.

Potassium hydroxide (KOH) is highly hygroscopic. Due to this feature, it is very easily soluble in water. The relationship of KOH to lipid structure is well known. Accidental use in humans has led to caustic injuries resulting in serious death due to cell membrane adipose tissue damage(8,9). In this study, the virucidal effect was aimed with the effect on the virus envelope by taking advantage of this bad reputation of KOH.

In previous studies on the antiviral properties of KOH, beneficial effects were demonstrated by buffering KOH to pH:8.9-9.0 in in vitro and in vivo studies(10). It has been reported in the previous study that optimization studies focused on pH:8.9-9.0 in KOH solution, as drinking water usage at pH:9.0 was observed in different locations around the world.

In the previous study with KOH, an exergonic reaction of KOH to the cell envelope phospholipid structure was observed in silico, however, it was observed that it increased the life span in cell culture(10). The fact that histopathological findings were not observed with KOH inhalation administered to mice in in vivo studies and that oxidative stress parameters were found to be more positive support the safety of KOH inhalation with this dosage(10).

Tomato spotted wilt virus (TSWV) is an enveloped RNA virus belonging to the genus Tospovirus in the Bunyaviridae family(11). TSWV causes epidemics in some areas. In these epidemics, many features such as plant species, variety and development period of the plant, type and density of virus carrier population, and ability to carry the virus have an important place (12).

As with any viral disease, symptoms begin to appear after a certain time after TSWV enters the host. In TSWV infection, symptoms such as mosaic, ring spots, yellowing, chlorotic lines, necrosis and deformation are observed on the leaves of plants (12,13).

In this study, to demonstrate the virucidal activity of KOH by destroying the lipid envelope structure of enveloped viruses; TSWV was used as the virus and pepper was used as the host test plant.

The study protocol is in the pattern below.

1. In silico detection of intermolecular interactions between different molecular numbers of KOH and molecules in the cell envelope.
2. Obtaining virus from plants infected with TSWV.
3. Inoculation of TSWVs interacted with different KOH concentrations into groups of healthy plants and symptomatological observation of plants.
4. Investigation of the presence of virus in the tissues of all plants, after observing the symptoms of the disease.
5. In silico studies, discussion of the results obtained after the interaction of KOH solution and TSWV.
6. Integrating these findings with previous studies with KOH and discussing the virucidal efficacy of KOH inhalation in the treatment of respiratory system enveloped virus infections.

## Material and method

### Ligand and Receptor retrieval

Human membrane bilayer complex with matrix metalloproteinase-12 used as the target receptor was downloaded from the RCSB PDB database (https://www.rcsb.org/) with the code 2mlr (resolution: not applicable). The matrix metalloproteinase component and all the heteroatoms surrounding the phospholipid bilayer complex were removed prior to docking. On the other hand, potassium hydroxide (KOH) used as the ligand against the phospholipid bilayer complex was drawn using GaussView 6.0.16 embedded in Gaussian 09 program and optimized with the B3LYP hybrid function (6-311G (d, p) basis set) (14).

In this molecular docking study of KOH against the human phospholipid bilayer complex (2mlr), the following hypothesis was established: Due to its highly reactive chemical structure, the KOH molecule may enter into favorable exergonic interactions (mostly disrupting) with the components forming the cell membrane and can establish non-bonded contacts with the hydrophilic (ex. choline, phosphate or glycerol) groups in the phospholipid bilayer. Indeed, docking calculations may not reveal the direct damage caused by KOH on these groups, however, the strength of the molecular affinity of KOH towards these regions can reliably be calculated. In addition to the single-ligand docking protocol, a simultaneous multiple-ligand docking protocol was also applied to observe the extent to which the binding strength (ΔG°=kcal/mol) changes with the increase in the number of KOH molecules on the phospholipid bilayer, and thus the synergistic effects of KOH molecules on the components of the cellular membrane were also shown. The simultaneous docking protocol is a fairly new approach in molecular docking field, thus, our study also provides new information to the literature on this aspect.

### Molecular docking between the KOH and human membrane bilayer complex using single and simultaneous docking protocols

Prior to docking simulations with AutoDock Vina, target (2mlr) and ligand (KOH) structures were prepared using AutoDockTools 1.5.6 and saved in pdbqt format (15,16). In molecular docking simulations with human phospholipid bilayer complex and KOH, polar hydrogen atoms in receptor and ligand molecule were retained, whereas, non-polar hydrogens were merged. Kollmann charges were assigned to the receptors and Gasteiger charges were assigned to the KOH.

In both the single- and multiple-ligand docking protocols, the same size of grid box with 50×50×30 Å points (x = 29.49; y = 31.39; z = 70.99) and 0.375 Å grid spacing was defined. This grid box size were adjusted in a way that the KOH molecule could easily interact with various components of the phospholipid bilayer complex.

After two separate (single and multiple) docking runs of KOH against the membrane phospholipid bilayer complex (20 independent dockings for each), all potential binding modes of the ligand were clustered by AutoDock Vina on the basis of geometrical similarity and were ranked based on the binding free energy (ΔG°; kcal/mol) of each of the ligand conformations which showed the lowest (best) affinity against the target receptor. The top-ranked docking conformations (modes) of the KOH molecule calculated by AutoDock Vina were rendered and analyzed using Discovery Studio Visualizer v16 (17).

### Preparation of KOH solution

KOH (Merck, Darmstadt, Germany) was added to distilled water at 25 degrees, and the solution was buffered in a magnetic stirrer (0,13951 mg/1 L) to pH 11. When the pH was fixed at 11, the process was continued with a magnetic stirrer for 5 minutes. Then, the solution was kept in an airtight glass container until the plants were treated.

### Mechanical Inoculation Studies

Mechanical inoculation studies were carried out at Çukurova University Faculty of Agriculture, Department of Plant Protection. Accordingly, plant tissues taken from a pepper plant, which was determined to be infected with TSWV by DAS-ELISA method, were mixed with phosphate buffer solution (0.1M, pH:7.0, 0.01M mercaptoethanol, 0.2% sodium sulfide) in a sterile mortar and 1/10 (g/ml)) was extracted by diluting (18). Then, different ratios of KOH solution (1 unit extract/1 unit KOH solution; 1 unit extract/3 unit KOH solution and 1 unit extract/7 units KOH solution) were added into the extract and inoculated on the leaves of the test plants that had previously been sprinkled with carborandum powder. After the application, the test plants were kept in the air-conditioning room at 22-24 °C temperature, 70% humidity, 16/8 hours (Light/Dark) and 5000 Lux lighting conditions in order to observe the symptom output. Inoculated plants were checked periodically every day until the 45th day of the study for the symptoms of mosaic, ring spots, yellowing, chlorotic lines, necrosis and deformation on the leaves.

### Serological Studies

Test plants inoculated with TSWV in mechanical inoculation studies were tested by DAS-ELISA method using TSWV-specific ELISA kit (BIOREBA Art.Nr:19075) on the 30th and 45th days after inoculation. DAS-ELISA tests were performed according to the method recommended by the company where the ELISA kit was determined. One gram of the newly formed leaf tissues of pepper plants was used in the tests.

Accordingly, 100 µl of each sample prepared in sample buffer solution was added to the wells of the ELISA plate (NUNC) coated with TSWV-specific γ-globulin and incubated for 16 hours at +4 °C. After the washing step, conjugate was added to the wells and the plates were incubated at 35 °C for 4 hours. As the last step, the prepared substrate was added at a concentration of 1mg/ml in the substrate buffer solution and readings were started after about 60 minutes.

The results were read on the Thermo MULTISKAN GO reader at a wavelength of 405 nm. Two controls, negative and positive control, were used in each ELISA plate, and samples that gave absorbance values at least twice the value read from the negative control as a result of the tests were considered positive (19). The results were evaluated by averaging the absorbance values of the wells used for each sample.

### Statistical evaluation

Non-parametric MannWhitney U test was used to compare the means of two independent groups that did not show normal distribution characteristics (for evaluation of ELISA and symptomatological findings). SPSS 17 package program was used in the study. p<0.05was considered significant.

## Results

### Interactions of potassium hydroxide (KOH) with human membrane bilayer complex

We conducted molecular docking simulations to demonstrate the intermolecular interactions of potassium hydroxide (KOH) with the components of human membrane bilayer complex (2mlr). In addition, the molecular affinities (binding free energy [^Δ^G°]; kcal/mol) of both single and multiple KOH molecules against the phospholipid membrane were also determined.

Considering the single KOH docking results, it was determined that KOH formed a hydrogen (H) bond with the O7 atom of the glycerol, an attractive charge (electrostatic) interaction with the O1 atom of the phosphate as well as a metal-acceptor interaction with the O3 atom of the same phosphate on two adjacent phospholipid molecules of the membrane structure (Figure 1). The calculated binding affinity of the KOH molecule in this region of the membrane bilayer complex is energetically favorable (^Δ^G°= −2.21 kcal/mol) (Table **1**).

**Figure 1.**
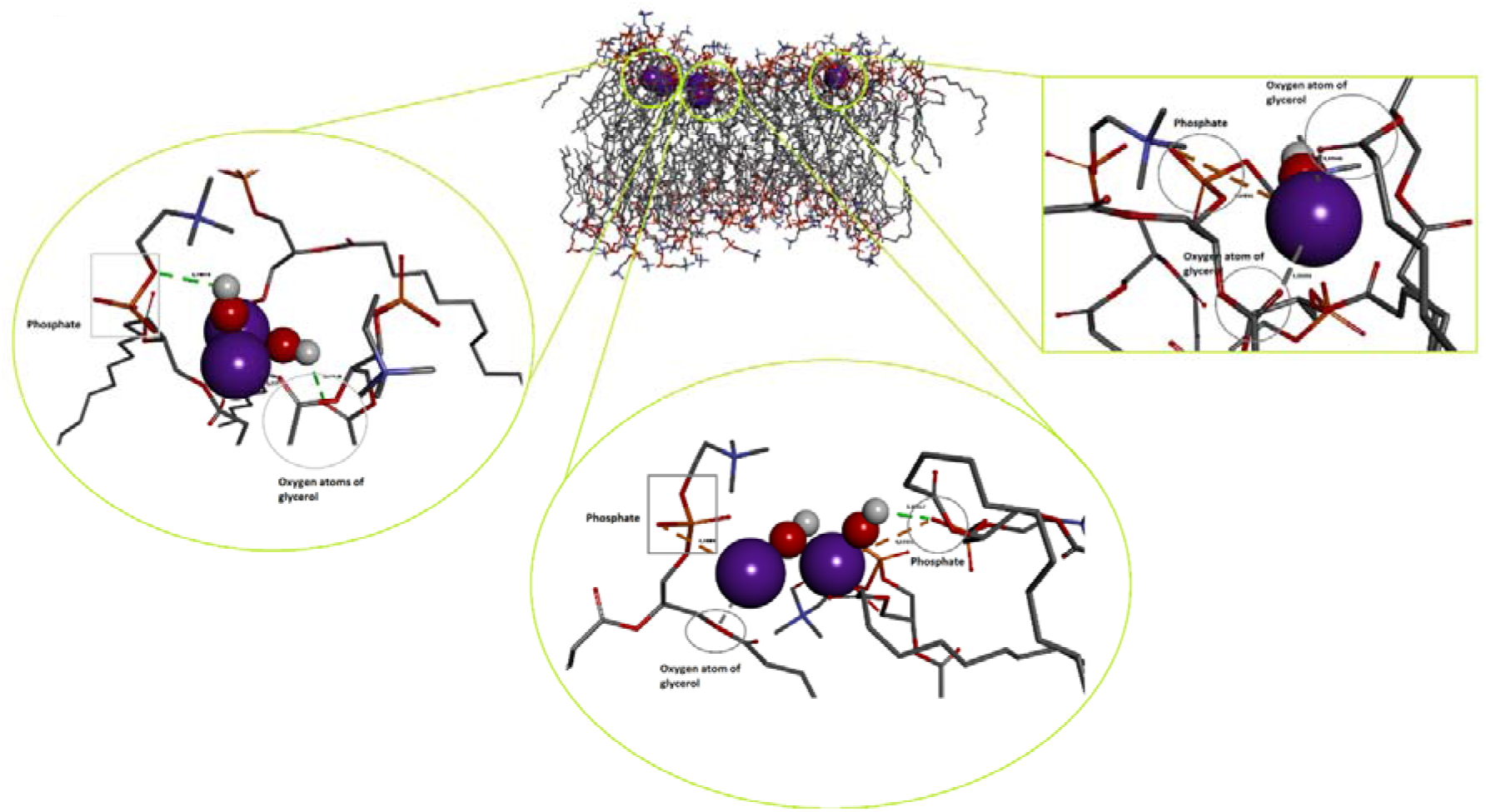
3D view of the docking interactions of multiple (5) potassium hydroxide (KOH) molecules with the human membrane bilayer complex (2MLR). Green dashed lines indicate hydrogen bonds, orange dashed lines show electrostatic interaction, and grey dashed lines depict metal-acceptor interaction. KOHs are shown in CPK mode and membrane bilayer complex is shown in the stick mode. Visualization and rendering was performed with DS Studio v16.

Docking of the multiple KOH molecules against the human membrane bilayer complex led us to achieve even more supportive and expected results. The simultaneously docked 5 molecules of KOH formed two H-bonds with the O2 and O3 atoms of phosphates and O6 atom of glycerol, electrostatic interactions with the O1 and O2 atoms of phosphates and metal-acceptor interactions with the O5 and O8 atoms of glycerol groups (Figure 1). The calculated binding affinity of the KOH molecules in those regions of the membrane bilayer complex was found to be energetically highly favorable (ΔG°= −10.83 kcal/mol) (Table 1).

**Table 1.** Predicted binding free energies and membrane bilayer complex (2MLR) interaction results for potassium hydroxide (KOH) docked in single and multiple forms.

| Compound | Receptor | $\Delta G_{\text{best}}$<br>(kcal/mol) | Classical<br>H-bond | Electrostatic | Other<br>(metal-acceptor) |
| --- | --- | --- | --- | --- | --- |
| *Single KOH* |  |  |  |  |  |
| KOH | 2MLR | -2.21 | Glycerol<br>(O7) | Phosphate<br>(O1) | Phosphate<br>(O3) |
| *Multiple KOH* |  |  |  |  |  |
| 5 KOHs | 2MLR | -10.83 | Phosphate<br>(O2, O3),<br>Glycerol<br>(O6) | Phosphate<br>(O1, O2) | Glycerol (O5, O8) |

$\Delta G_{\text{best}}$ : most favorable binding free energy (kcal/mol) 2MLR: Membrane bilayer complex with matrix metalloproteinase-12 at its alpha-face O: Oxygen atom

Molecular docking results indicate that the strength of the interaction of multiple KOH molecules with the membrane is considerably higher (ΔG°= −10.83 kcal/mol), and most likely their membrane-damaging reactivity is also significantly increased when compared to single KOH docking (ΔG°= − 2.21 kcal/mol). Indeed, these findings should be verified with appropriate biological or kinetic experiments.

### Symptomatological findings in plants

In the group inoculated only with TSWV for the control purpose, all of the plants had mosaic, ringed spots, yellowing, chlorotic lines, necrosis and deformation findings on the 30th and 45th days symptomatologically (Figure 2 D). No symptomatological findings were observed in the groups inoculated with TSWV interacting with KOH (p<0.01) (Figure 2 E,B).

**Figure 2:**
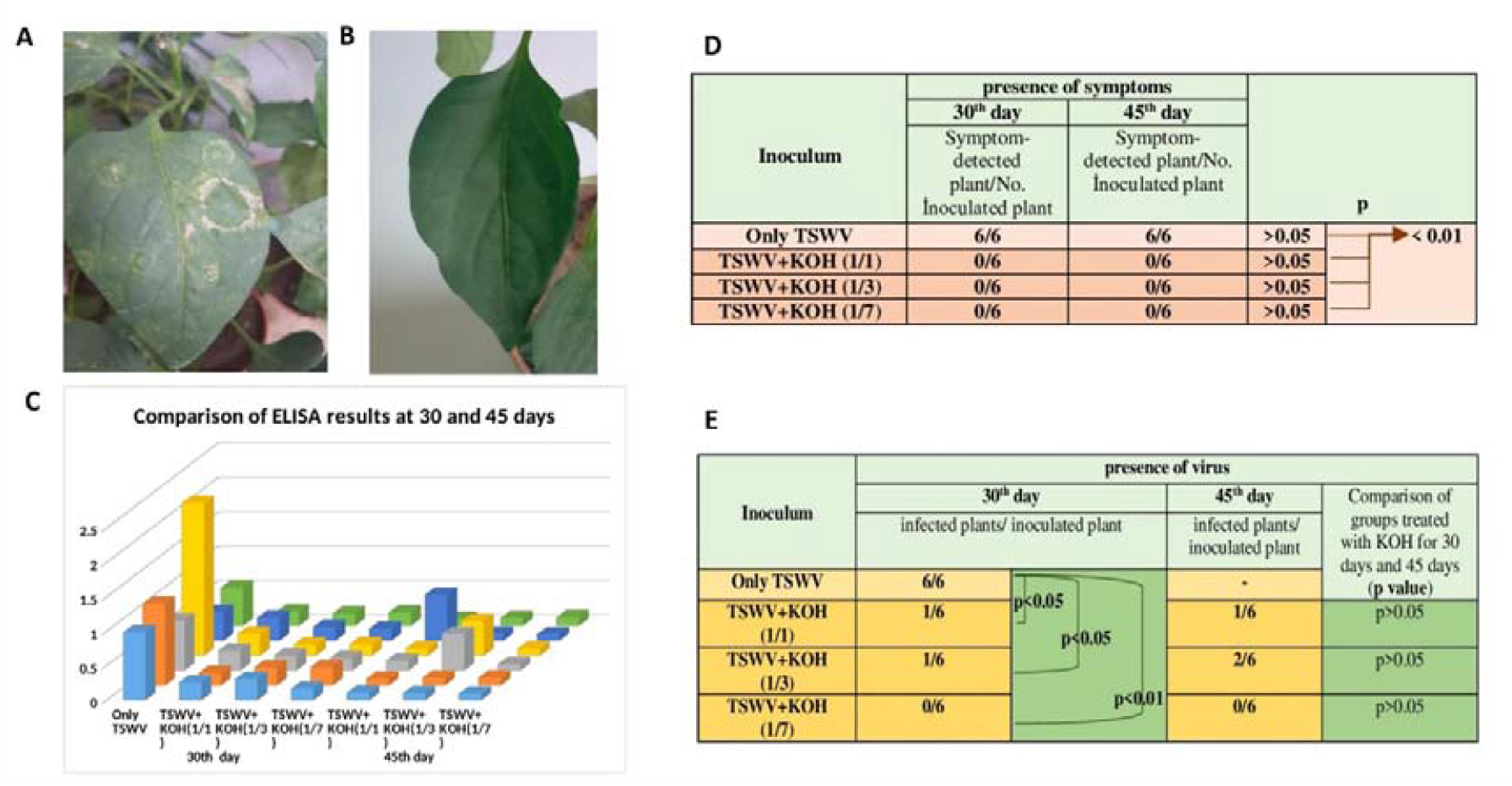
**A**. Plant inoculated with TSWV without KOH interaction. **B**. Plant inoculated with TSWV interacting with KOH. **C**. The average ELISA absorbance values of all plants individually (absorbance values are given in the supplement file). **D**. Presence of symptoms and statistical differences in groups. **E**. Presence of virus in the groups and statistical evaluations between groups, differences between the 30th and 45th days.

### Detection of the presence of viruses in plants

DAS-ELISA results were found positive in all control plants (6/6) inoculated only with TSWV. In the groups inoculated with viruses interacting with KOH, positive absorbance value was detected in 1 plant (1/6) (p<0.05) with 1/1 interaction (1 mL extract+1 mL KOH solution), and positive absorbance value was detected in 1 plant (1/6) with 1/7 interaction (p<0.05) (1 mL extract+7 mL KOH solution). No positive absorbance value was detected in any plant in the 1/3 interacting group (p<0.01) (1 mL extract+3 mL KOH solution) (Figure 2 C,E).

## Discussion

It seems very difficult to apply clinical studies to evaluate the therapeutic agent efficacy in viral diseases. In this study, where we focused on observing the virucidal activity of KOH in enveloped viruses, we conducted in silico studies. Then we focused on the biological equivalent of these simulation studies.

In the previous simulation study with KOH, the intermolecular interaction of KOH in the lipid envelope structure was found to be −1,62 kcal/mol(9). In our study, a single KOH interaction with glycerophosphate molecules in the lipid envelope layer was found to be −2.21 kcal/mol. This data was in agreement with the previous result. However, in our study, it was observed that the exergonic energy effect increased up to −10.83 kcal/mol with the increase of KOH molecules in the medium (Table-1).

Glycerophospholipids are the primary building block of the cell membrane. The two fatty acids formed ester bonds with the first and second hydroxyl groups of L-glycerol-3-phosphate. The ester bonds formed are one of the basic structures in maintaining the integrity of the cell membrane (20).

In the study, 3 different interaction points were found with the lipid envelope of KOH. These are 1) the bond with the hydrophilic R group attached to the phosphate on the envelope, 2) the bond with the phosphate below the R group, and 3) the ester bonds between glycerol and the fatty acid chain (Figure 3B).

**Figure 3:**
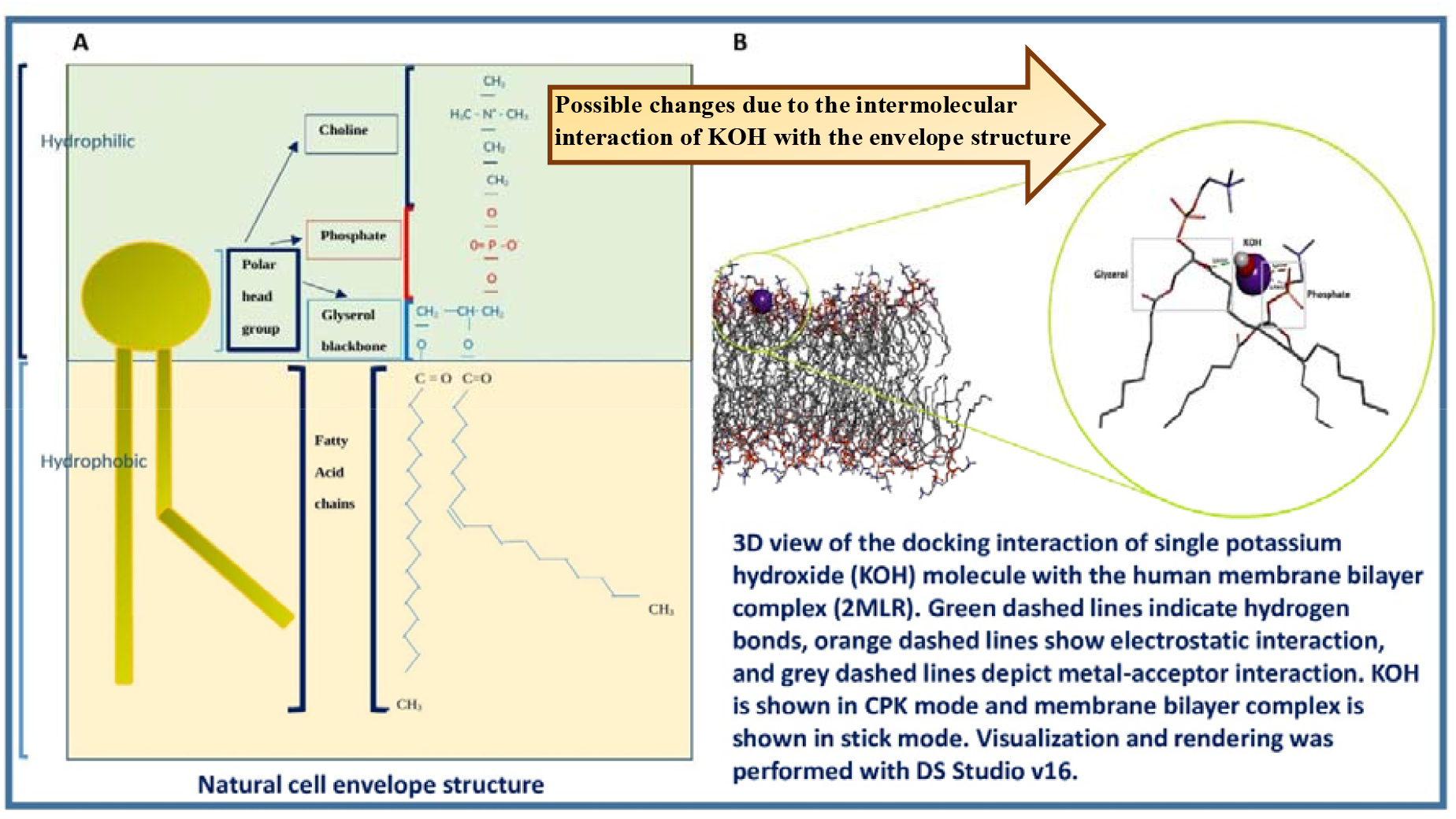
**A**. Hydrophilic and hydrophobic parts of the normal cell envelope and important bonds of the glycerol backbone. **B**. Simulation of possible changes in bond structures with the addition of KOH to the medium.

If even a small amount of water is present, significant hydrolysis of lipids to free fatty acids with KOH may occur, depending on the reaction length(21,22). In the light of this information and results, we think that KOH destroys the envelope lipid layer with nucleophilic attack on the bonds it interacts with, especially the ester bonds, since it is very reactive.

In-tube suspension test was not performed to evaluate virucidal activity. We thought that false positive results could be detected in the measurement of RNA released as a result of envelope destruction as a result of the interaction of the virus envelope with KOH(23). For this purpose, we wanted to evaluate a biological process in which we can observe the infectivity of the virus. For this, we chose TSWV as the virus and the test plant pepper seedling as the virus host.

TSWV has the ability to cause disease in pepper. It creates the disease by entering from injured areas on the outer surface of plants (24). For this purpose, virus inoculation was facilitated by destroying the leaves with carboronium. After TSWV enters the plant from the damaged area, it spreads in the plant and causes disease. In our study, all the symptoms were present on the 30th day in all plants in the group inoculated with viruses that did not interact with KOH. Serological detection of TSWV in all plants in the same group on the 30th day shows us the reliability of the technique of implantation of the virus into the tissue. In the groups with KOH and virus interaction, TSWV was detected in 1/6, 0/6 and 1/6 plants, respectively (Figure 2C,E), the difference in the findings was statistically significant (p<0.05). In accordance with this, the presence of 1/6, 0/6 and 2/6 viruses, respectively, in the groups where no statistical difference was observed on the 45th day in plants inoculated with TSWV interacting with KOH increases the reliability of the study (p>0.05).

Exergonic reactions of intermolecular interactions in the lipid envelope structure, KOH’s affinity for ester bonds between glycerol and fatty acid chain, and the fact that all reactions take place in environments containing water strengthen the nucleophilic attack of KOH. These results obtained in the study show that KOH destroys the envelope by making lipid hydrolysis especially with glycerophosphate interaction in the envelope structure of TSW viruses and eliminates virus infectivity. Since the aim of this study was to show the destruction of lipids in the envelope structure upon contact of KOH with an enveloped virus, the information obtained in the discussion was discussed together with viruses that cause disease in the respiratory tract, epidemics and pandemics.

However, it is known that KOH is a highly corrosive substance. Does the use of this inhaler containing corrosive molecules cause damage to the lungs?

Hypochlorous acid (HOCl), which the body produces and uses itself, can be a good example of this. HOCl is a highly irritating molecule for humans. However, HOCl is used in body defense by being formed from H2O2, Cl and H by myeloperoxidase in leukocytes(25,26). If HOCl is not used, many diseases that can result in death can be encountered. But the HOCl used in the cell is at a very low concentration. Therefore, it can be thought that it can be used in treatment by regulating the KOH concentration with optimization studies.

For this reason, in the previous study with KOH, the highest pH values of the water used as drinking water were taken as the upper limit in order to prevent the alkaline irritant effect of KOH. This value is pH: 8.9-9.0. In the transportation of KOH to the lungs by inhalation, distilled water was supported by a NaCl concentration compatible with the interstitial fluid (0.89-0.9%)(10).

In addition to the interactions of KOH in the cell envelope structure, the observed findings support the safety of KOH inhalation. These findings; It is the exergonic intermolecular interaction of KOH to the coronavirus spike glycoprotein active site, human AGE2 active site, H. influenza virus neurominidase enzyme active site. In addition, it was determined that in cell culture (MTT) fibroblast survival time was extended by 49% in 24 hours. In a study in mice, it was found that KOH oral spray and inhalation did not cause histopathological findings in the oral region and lungs, and in addition, it positively changed oxidative stress markers(10).

The virus is much smaller than a human cell (Figure 4). Therefore, the virus envelope will be much easier to contact and be affected by KOH molecules than the human cell envelope. In addition, the antioxidant defense system, which is not in the virus but in the human cells, also turns this paradoxical activity of KOH in favor of the human cell.

**Figure 4:**
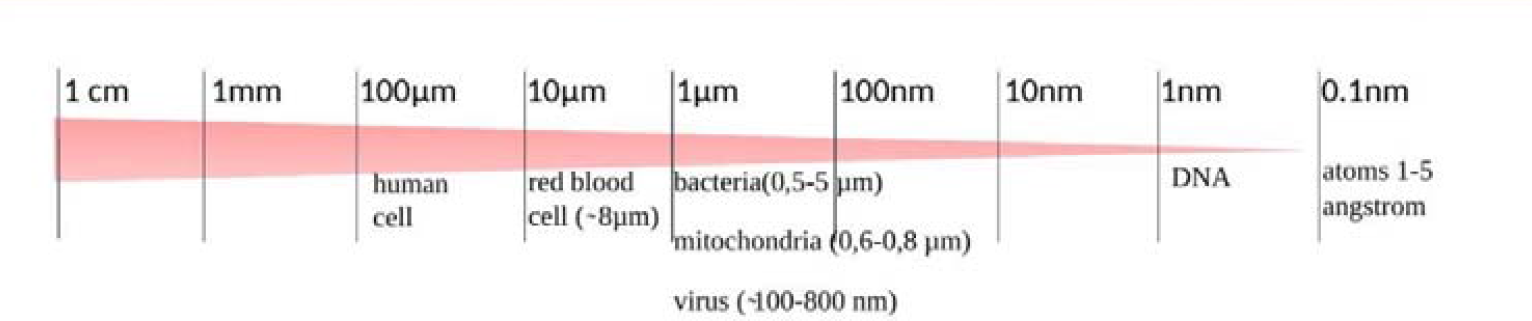
Comparison of virus size with human cell size.

In a non-invasive clinical study with KOH, it was observed that the interaction of KOH in human mucus reduces the surface tension and contact angle, and shifts the mucus to the alkaline side. It has been determined that KOH inhalation in mice reduces the contact contact angle in bronchoalveolar lavage. These results emphasize the mucolytic effect of KOH as stated in the study. In these, it contributes positively to the virucidal activity of KOH in the treatment(10).

In another study with the KOH molecule, the affinity of KOH for disulfide bonds in some human immunoglobulins (Ig) Fab region was demonstrated by intermolecular interactions (27). These findings suggest that KOH potentially induces low-weight Ig production. Low-weight Ig with high mobility in mucus whose fluidity decreases with inflammation will be very effective for opsonization. It will be effective in preventing diseases in early Ig response by recognizing antigens much more easily in virus mutations.

In enveloped viruses, the destruction of the virus envelope structure causes the genetic material in the virus to lose its function. As a result of this effect, the virus loses its infectivity (28). This feature, which is valid for all enveloped viruses, is valid for all enveloped viruses that cause disease in humans, animals and plants.

As a result, KOH showed antiviral effect in TSWV. According to the results of this study, it is thought that KOH makes this effect with the destruction of the virus envelope structure lipid (Figure 5).

**Figure 5:**
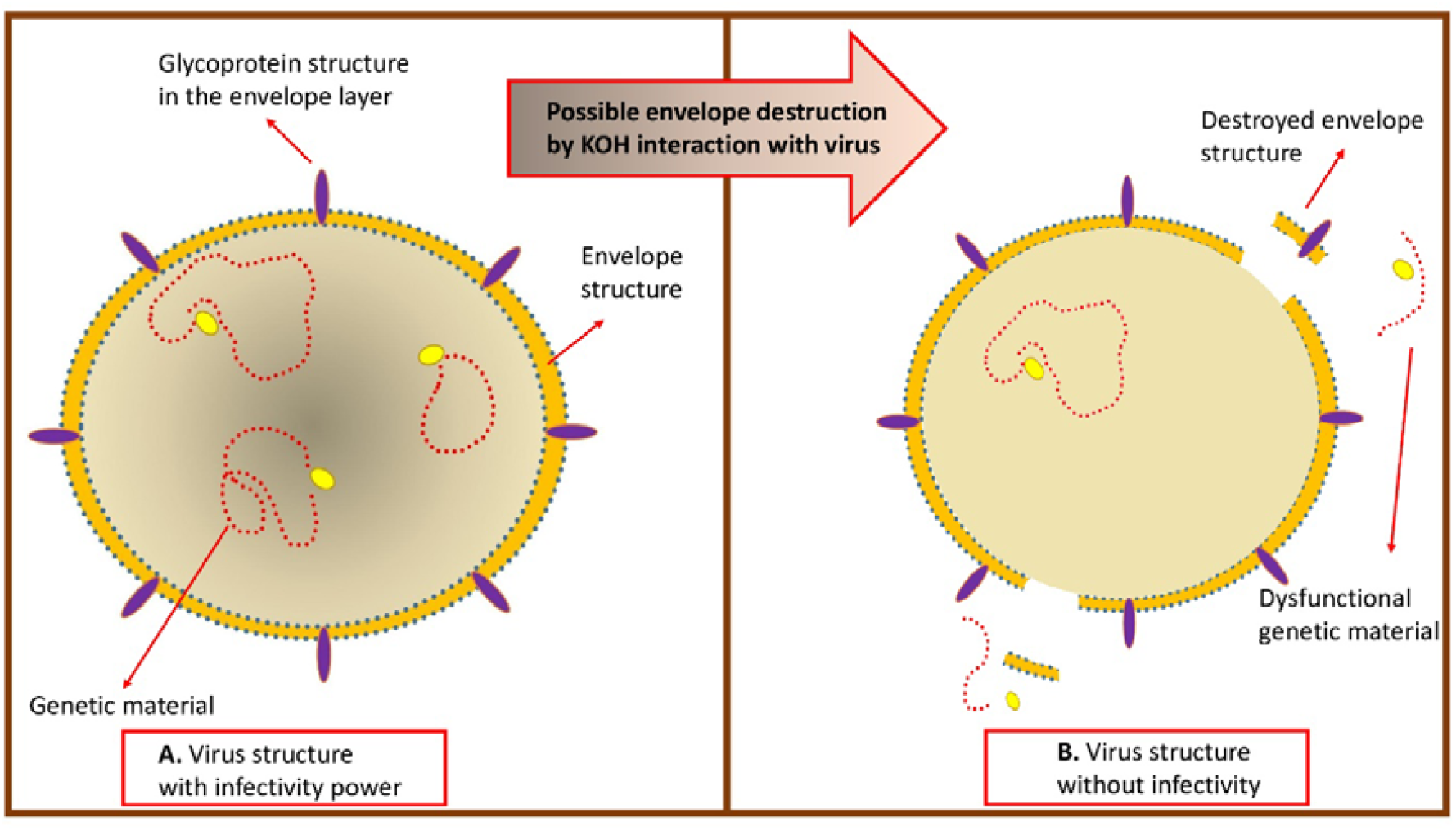
**A**. Virus structure with infectivity power, **B**. Virus structure without infectivity When the virucidal effect of KOH is added to the mucolytic, alkalinizing and small-weight Ig generation potential, enveloped viruses such as H. influenza, Coronavirus, Monkeypox, or mutant types of these viruses, which progress with epidemics and pandemics, can be treated with KOH inhalation.

Influencing (KOH) and affected (enveloped virus) agents were brought together for evidence-based scientific data at the common point of life. The effects created by KOH on the functioning so far on natural life were observed. The response of the plant, which was tried to be infected with the enveloped virus, but whose life was positively affected by the effect of KOH, to the virus attack was discussed. The data obtained by the multidisciplinary collaboration, the clinical status of the respiratory system cells infected with enveloped viruses and the effectiveness of KOH inhalation therapy were discussed.

This study makes us think about the positive efficacy of KOH inhalation on the fate of enveloped virus-infected alveoli in the respiratory system and thus human life. Clinical studies are needed for the therapeutic efficacy of KOH.

Although the interpretations of our study are for the treatment of enveloped viral diseases involving the respiratory system, further studies are needed with KOH in tomato spotted wilt virus disease, since there is still no antiviral drug to prevent TSWV.

There is no conflict of interest between the authors

## Supporting information

Supplement Figure-2, page-6

