## Supplement Figure-2, page-6 for "Virucidal Activity of Potassium Hydroxide Modeling on Tospovirus"

|  | **30^th^  day** | | | | **45^th^ day** | | |
| --- | --- | --- | --- | --- | --- | --- | --- |
| **Plant no** | **Only TSWV** | **TSWV+ KOH(1/1)** | **TSWV+ KOH(1/3)** | **TSWV+ KOH(1/7)** | **TSWV+ KOH(1/1)** | **TSWV+ KOH(1/3)** | **TSWV+ KOH(1/7)** |
| 1 | **0,987** | 0,258 | 0,306 | 0,167 | 0,1097 | 0,1019 | 0,0941 |
| 2 | **1,187** | 0,18 | 0,262 | 0,264 | 0,097 | 0,1101 | 0,1112 |
| 3 | **0,726** | 0,288 | 0,216 | 0,18 | 0,141 | 0,5271 | 0,0902 |
| 4 | **2,216** | 0,316 | 0,157 | 0,163 | 0,1016 | 0,5026 | 0,1022 |
| 5 | **0,417** | 0,334 | 0,199 | 0,174 | 0,6676 | 0,0996 | 0,0925 |
| 6 | **0,556** | 0,207 | 0,177 | 0,192 | 0,0988 | 0,1115 | 0,1232 |
| Negative control | 0,152 | 0,159 | 0,147 | 0,145 | 0,0929 | 0,0847 | 0,0907 |
| Positive control | 0,236 | 0,163 | 0,197 | 0,171 | 0,1453 | 0,2215 | 0,1571 |
